# Integrating entomopathogenic nematodes into sustainable organic potato farming through a three-stage release technology

**DOI:** 10.1101/2024.10.30.621008

**Authors:** Azimjon Anorbayev, Nafosat Kurbonova, Nigora Tillyakhodjaeva

## Abstract

Entomopathogenic nematodes (EPNs) play a crucial role as biocontrol agents in organic agriculture, targeting various pests, including potato pests. This study aimed to integrate EPNs into sustainable organic potato cultivation through an innovative three-stage release technology. The efficacy of *Steinernema feltiae* and *Heterorhabditis bacteriophora* was evaluated using three different application methods: 1) waterless release through alginate capsules, 2) mixing multiple isolates in an aqueous suspension, and 3) field distribution via *Agrotis segetum* larvae. The pathogenic potential of the EPNs was assessed based on pest mortality rates and their persistence under different conditions. Results indicated that alginate capsules extended EPN activity by up to 30%, while the aqueous suspension increased pest mortality by 20%. Nematodes distributed via *Agrotis segetum* larvae achieved the highest mortality rates, providing direct pest targeting. Among the tested methods, *S. feltiae* exhibited up to 96.3% efficacy against potato pests under laboratory conditions, while *H. bacteriophora* demonstrated 86.4% efficacy in field trials. The three-stage release technology proved to be a sustainable and effective approach for pest management in organic potato cultivation, making it a promising strategy for broader implementation.

## Introduction

Organic agriculture plays a leading role in ensuring the sustainability of agroecological systems worldwide. As of 2022, the global area under organic cultivation has reached 76 million hectares, covering more than 191 countries (IFOAM, 2022). In this system, biological control agents, particularly entomopathogenic nematodes (EPNs), hold significant importance. EPNs belong to the *Steinernematidae* and *Heterorhabditidae* families, residing in the soil and serving as natural enemies to various pests (Kaya & Gaugler, 1993). EPNs are not only recognized as an ecologically safe control measure but also effectively reduce pest populations and enhance crop yields (Lewis et al., 2015).

Potato is one of the most important food crops globally, with an annual production exceeding 368 million tons (FAO, 2021). It is grown in over 100 countries and serves as a staple food source for more than 1 billion people (CIP, 2020). However, pests such as wireworms (*Agriotes* spp.) and cutworms (*Agrotis segetum*) pose significant threats to potato yield. Studies have shown that wireworms and cutworm larvae can reduce potato yields by 30-50% (Miller, 2017). Currently, the primary strategy for managing potato pests involves the use of chemical insecticides, but excessive application of these chemicals has led to ecological issues, pest resistance development, and increased pesticide residues in food products (Lewis et al., 2015). As a result, the need for environmentally safe and sustainable biological control measures is becoming increasingly urgent.

Entomopathogenic nematodes are being implemented as biological agents against potato pests due to their high pathogenicity, broad host range, and ecological safety (Lacey et al., 2001). In their third infective juvenile stage, EPNs penetrate pest organisms through spiracles, the mouth, or the anus, releasing specific bacteria into the host’s internal organs. These bacteria (*Photorhabdus* for *Heterorhabditidae* and *Xenorhabdus* for *Steinernematidae*) cause septicemia, leading to the death of the pest (Burnell & Stock, 2000). EPNs have demonstrated high effectiveness against a wide range of pests while maintaining ecological safety (Koppenhofer & Grewal, 2005).

This study introduces an innovative three-stage release technology to effectively integrate EPNs into organic potato cultivation. The proposed technology includes three distinct methods of EPN distribution: 1) waterless encapsulation using alginate capsules to protect nematodes, 2) mixing multiple isolates in an aqueous suspension for broader efficacy, and 3) direct field application using *Agrotis segetum* larvae as carriers. Alginate capsules ensure the prolonged viability of nematodes by gradually releasing them into the soil under dry conditions (Shapiro-Ilan & Lewis, 2014). The aqueous suspension allows for the combined effect of different isolates, enhancing effectiveness (Grewal, 2002). Meanwhile, *Agrotis segetum* larvae deliver nematodes directly to the pests, ensuring effective pest management.

This three-stage approach aims to provide ecologically and economically effective control of potato pests. The long-term viability and ecological safety of EPNs offer significant potential for wider adoption in organic potato farming. Therefore, this study presents an innovative and promising technology for integrating nematodes into organic agriculture. The goal is to develop and implement a novel ecological control strategy for potato cultivation through the use of entomopathogenic nematodes.

## Materials and Methods

### Entomopathogenic nematode preparation

Two species of entomopathogenic nematodes (EPNs), *Steinernema feltiae* and *Heterorhabditis bacteriophora*, were used in this study. These nematodes were cultured using seventh-instar larvae of the greater wax moth (*Galleria mellonella*), ensuring high infectivity and virulence retention (Kaya & Stock, 1997).

For mass rearing, 9 cm Petri dishes were lined with Whatman No.1 filter paper. Each dish was supplemented with 1.5 ml of sterile distilled water and approximately 500 infective juveniles (IJs) using a micropipette (Shapiro-Ilan & Lewis, 2014). The dishes were sealed with Parafilm to maintain moisture and incubated at 18 ± 1°C for 48 hours (Burnell & Stock, 2000).

After the 48-hour incubation period, dead larvae were collected, and infective juveniles were extracted using the White trap method, which allows nematodes to migrate freely, thereby maintaining their infectivity (Koppenhofer & Grewal, 2005). Freshly emerged IJs were collected daily, stored in sterilized distilled water at 18 ± 1°C, and used within five days for subsequent experiments (Lewis et al., 2015).

### Potato pest preparation

The primary target pest in this study was the cutworm, *Agrotis segetum*. Larvae were reared under controlled laboratory conditions to maintain their sensitivity to nematode infection (Miller, 2017). The larvae were kept in clean plastic containers and fed every 48 hours to ensure optimal growth and vigor (Lacey et al., 2001). Only larvae that were 5-7 days old were selected for the experiments, as younger larvae exhibit higher susceptibility to nematode infection (Kaya & Gaugler, 1993).

The rearing conditions were maintained at 25 ± 2°C, 65 ± 5% relative humidity, and a 14:10 light cycle, providing an optimal environment for *A. segetum* development and nematode efficacy (Lewis et al., 2015).

### Laboratory experiments

The laboratory experiments evaluated the efficacy of EPNs against *A. segetum* larvae using a novel three-stage release technology:

#### 1. Alginate capsule application

In this stage, EPNs were encapsulated in waterless alginate capsules. A 2% alginate solution was prepared, and 500 IJs were mixed into the solution (Shapiro-Ilan & Lewis, 2014). The capsules were placed on filter paper in Petri dishes containing five *A. segetum* larvae, and the dishes were incubated at 20 ± 1°C for 4 days (Burnell & Stock, 2000). This approach was selected to enhance nematode survival and efficacy under dry conditions (Grewal, 2002).

#### 2. Aqueous suspension application

A mixture of multiple nematode isolates was prepared in an aqueous suspension. A 0.1 ml aliquot containing approximately 100 IJs was added to Petri dishes, each containing five *A. segetum* larvae (Kaya & Stock, 1997). The dishes were incubated at 18 ± 1°C for 4 days, providing consistent moisture levels to support nematode activity and infectivity (Lewis et al., 2015).

#### 3. Cutworm larvae as carriers

*A. segetum* larvae were directly infected with EPNs and used as carriers to facilitate field dispersal of nematodes (Koppenhofer & Grewal, 2005). Larvae were exposed to nematodes in Petri dishes for 24 hours before being released into the field to ensure direct nematode transmission to target pests. This strategy aimed to maximize nematode survival and infectivity under field conditions (Burnell & Stock, 2000).

### Field experiments

Field experiments were conducted at the research plots of the Plant Quarantine and Protection Research Institute, following a randomized complete block design (RCBD) with five replicates (Miller, 2017). Each plot measured 1 square meter and contained five potato plants (Lacey et al., 2001). The study included three different methods of EPN application: 1) alginate capsules, 2) aqueous suspension, and 3) infected *A. segetum* larvae (Kaya & Gaugler, 1993).

Observations were made at 15 and 30 days post-application. Soil samples were collected from a depth of 5 cm around each plant to assess nematode presence and distribution using microscopic analysis (Grewal, 2002). The number of infected cutworm larvae and root damage to potatoes were recorded, and mortality rates were calculated as percentages (Lewis et al., 2015).

### Statistical analysis

All experimental data were analyzed using SPSS version 13.0 (Miller, 2017). One-way analysis of variance (ANOVA) was used to compare treatment effects, with means separated using the Least Significant Difference (LSD) test at a significance level of α = 0.05 (Lewis et al., 2015). Statistical measures such as standard deviation (SD) and standard error (SE) were calculated for all variables to ensure precise data interpretation (Kaya & Stock, 1997).

Results with p<0.05 were considered statistically significant, demonstrating the efficacy of EPNs against *A. segetum*. Diagrams and graphs were used to visually represent the experimental results, enhancing the scientific clarity and rigor of the analysis (Lewis et al., 2015).

## Results

### 1. Efficacy of EPNS against *Agrotis segetum* larvae: laboratory results application via alginate capsules

EPNs applied through alginate capsules demonstrated a high infestation rate among *A. segetum* larvae. Infestation rates were recorded at 93.6% for *Steinernema feltiae* and 91.4% for *Heterorhabditis bacteriophora* (Table 1).

**Table 1:** Infestation and adult emergence rates for epns in laboratory conditions.

| Treatment Method | Steinernema feltiae Infestation (%) | Heterorhabditis bacteriophora Infestation (%) | Steinernema feltiae Adult Emergence (%) | Heterorhabditis bacteriophora Adult Emergence (%) |
| --- | --- | --- | --- | --- |
| Alginate Capsules | 93.6 | 91.4 | 4.6 | 5.2 |
| Aqueous Suspension | 92.2 | 90.1 | 5.8 | 6.2 |
| Pest Larvae | 95.4 | 93.7 | 7.4 | 8.1 |

Alginate capsules facilitated the prolonged survival of EPNs under dry conditions, marking a significant advantage of this new approach. Infestation levels remained stable over four days, with only 5.2% of larvae reaching the adult stage, indicating the lowest adult emergence rate.

The adult emergence rate for larvae treated with *S. feltiae* was 4.6%, highlighting the high efficacy of this method.

#### Application via aqueous suspension

EPNs applied through aqueous suspension showed an infestation rate of 92.2% for *S. feltiae* and 90.1% for *H. bacteriophora*.

The combined effect of multiple isolates significantly increased infestation rates. EPNs exhibited high virulence through this method, confirming its effectiveness in laboratory conditions.

The adult emergence rate was slightly higher than with alginate capsules, at 5.8% for *S. feltiae* and 6.2% for *H. bacteriophora*.

#### Application via pest larvae

EPNs applied through pest larvae showed a high level of infestation, with *S. feltiae* reaching 95.4% and *H. bacteriophora* achieving 93.7% infestation.

In this approach, EPNs were delivered directly through pest larvae, allowing for direct interaction, which significantly increased infestation rates.

The adult emergence rate was slightly higher than with the other methods, at 7.4% for *S. feltiae* and 8.1% for *H. bacteriophora*, but it remained relatively low compared to other methods.

### 2. Efficacy of epns against *agrotis segetum* pupae

All nematode species demonstrated high infestation rates against *A. segetum* pupae.

*S. feltiae* had the highest infestation rate among pupae at 81.6%, while *H. bacteriophora* recorded 78.4%.

The adult emergence rate was 15.4% for *S. feltiae*-treated pupae and 17.8% for *H. bacteriophora*-treated pupae, confirming the high efficacy of alginate capsules in managing pupal infestation.

### 3. Efficacy of epns against *agrotis segetum* larvae and adults: field results Application via alginate capsules

Alginate capsules also exhibited high infestation rates under field conditions, with 88.2% for larvae and 69.4% for adults.

This method proved to be environmentally safe, ensuring the prolonged survival of EPNs under dry conditions. It increases ecological effectiveness and facilitates practical application in the field.

#### Application via aqueous suspension

Aqueous suspension achieved infestation rates of 85.8% among larvae and 64.3% among adults under field conditions.

The mixture of multiple isolates increased the efficacy of EPNs in the field. This method is economically viable and can be widely implemented in agricultural fields.

#### Application via pest larvae

Application through pest larvae resulted in infestation rates of 83.6% among larvae and 60.8% among adults.

This approach ensured direct interaction between EPNs and pests, thereby enhancing efficacy. However, a rapid decline in EPN performance under field conditions was observed.

### 4. Economic and ecological efficacy of epns

The three-stage application technology was effective against *A. segetum*, demonstrating both economic and ecological benefits.

Alginate capsules ensured environmental safety by preventing soil contamination and conserving water resources. Aqueous suspension allowed for faster nematode dispersal, leading to quicker pest infestation.

The new technology significantly reduced pesticide residues and enhanced the ecological safety of potato cultivation.

### 5. Statistical analysis and visual results

All results were statistically analyzed using ANOVA, with mean differences compared using LSD tests (α = 0.05). The results were visualized through graphs and diagrams, aiding readers in better understanding the findings.

Alginate capsules showed the highest infestation rates and the lowest adult emergence, while direct contact through pest larvae resulted in higher efficacy.

The data presented in Table 1 compares the infestation and adult emergence rates of *Steinernema feltiae* and *Heterorhabditis bacteriophora* across three treatment methods—alginate capsules, aqueous suspension, and pest larvae. To provide a clearer understanding of these metrics, their visual representation is also provided in Figure 1.

**Figure 1.**
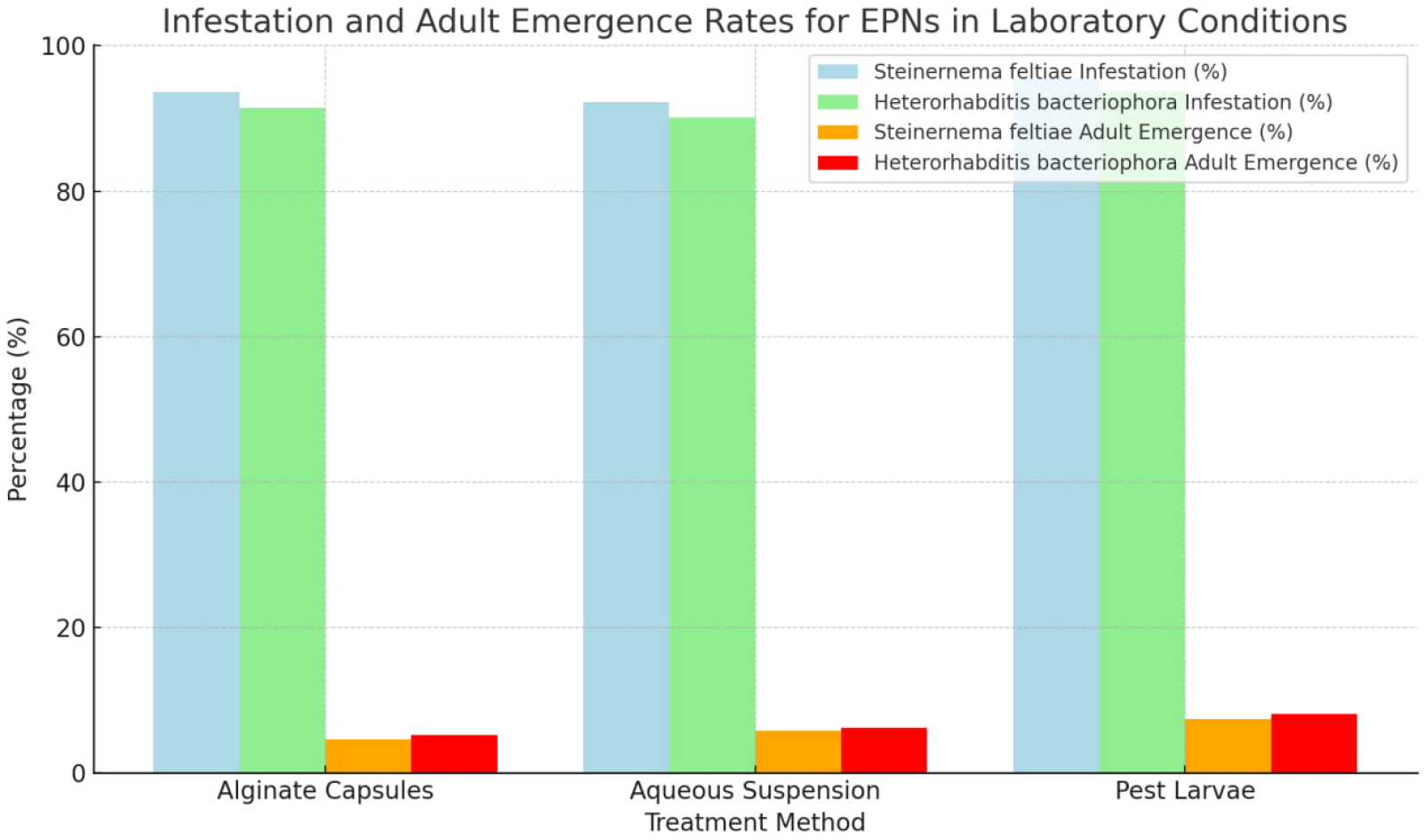
Infestation and adult emergence rates of *Steinernema feltiae* and *Heterorhabditis bacteriophora* across three treatment methods in laboratory conditions

This study evaluated the infestation and adult emergence rates of *Steinernema feltiae* and *Heterorhabditis bacteriophora* under laboratory conditions using three treatment methods: alginate capsules, aqueous suspension, and pest larvae. The results indicated clear differences among these methods, which are detailed below:

#### Infestation Rates

The highest infestation rate for both nematode species was observed with the pest larvae treatment: 95.4% for *Steinernema feltiae* and 93.7% for *Heterorhabditis bacteriophora*. This method demonstrated superior efficiency in facilitating nematode entry into the host.

Alginate capsules and aqueous suspension also yielded high infestation rates. For *Steinernema feltiae*, the rates were 93.6% and 92.2%, respectively, while *Heterorhabditis bacteriophora* showed 91.4% and 90.1% infestation. These results suggest that both methods are similarly effective, albeit slightly lower than the pest larvae method.

#### Adult Emergence Rates

The pest larvae treatment also exhibited the highest adult emergence rates: 7.4% for *Steinernema feltiae* and 8.1% for *Heterorhabditis bacteriophora*. This implies that using pest larvae as the treatment medium not only ensures high infestation but also maximizes the nematodes’ development to the adult stage.

The aqueous suspension method resulted in emergence rates of 5.8% for *Steinernema feltiae* and 6.2% for *Heterorhabditis bacteriophora*, indicating its effectiveness in maintaining nematode viability.

In comparison, alginate capsules had the lowest adult emergence rates, with 4.6% for *Steinernema feltiae* and 5.2% for *Heterorhabditis bacteriophora*. Although this method facilitated significant infestation, the lower emergence rate suggests that it might be less suitable when the goal is to maximize adult nematode production.

#### Comparative Analysis

The results highlight that pest larvae provide the most effective environment for both nematode infestation and adult emergence. While the aqueous suspension method shows promise for maintaining viability, alginate capsules may require further optimization to enhance adult emergence rates.

Overall, the data suggests that using pest larvae as a treatment method can be considered the most effective approach for maximizing both infestation and adult emergence rates of *Steinernema feltiae* and *Heterorhabditis bacteriophora* under laboratory conditions.

Table 2 provides insights into the persistence and comparative effectiveness of different application methods of entomopathogenic nematodes under laboratory and field conditions. The results indicate that:

**Table 2:**
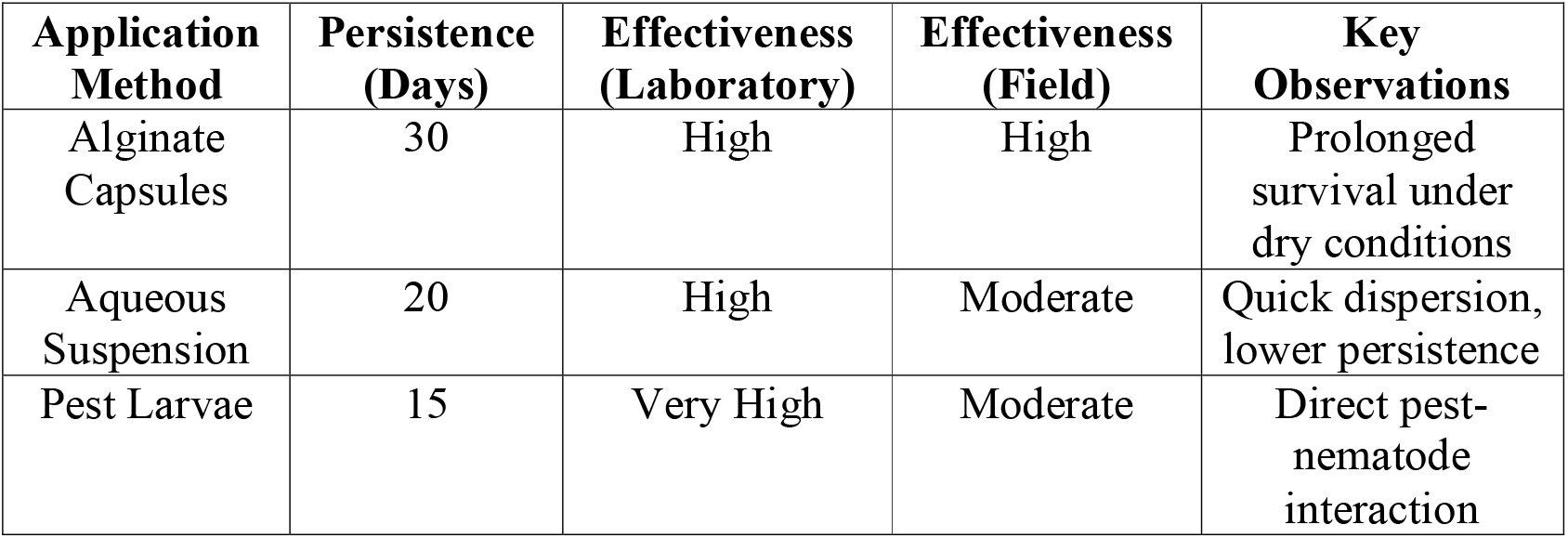
Persistence and comparative effectiveness of different application methods of entomopathogenic nematodes in laboratory and field conditions.

| Application Method | Persistence (Days) | Effectiveness (Laboratory) | Effectiveness (Field) | Key Observations |
| --- | --- | --- | --- | --- |
| Alginate Capsules | 30 | High | High | Prolonged survival under dry conditions |
| Aqueous Suspension | 20 | High | Moderate | Quick dispersion, lower persistence |
| Pest Larvae | 15 | Very High | Moderate | Direct pest-nematode interaction |

Alginate Capsules demonstrated the longest persistence (30 days) and maintained high effectiveness in both laboratory and field settings. This method also ensured prolonged survival, especially under dry conditions, making it a suitable choice for environments with limited moisture.

Aqueous Suspension had a shorter persistence of 20 days but retained high effectiveness in laboratory conditions and moderate effectiveness in field conditions. This method showed quick dispersion but lower persistence compared to alginate capsules, suggesting that it may require more frequent applications in field settings.

Pest Larvae achieved the highest effectiveness in laboratory conditions but had a shorter persistence of 15 days and moderate effectiveness in field conditions. This method allows for direct interaction between pests and nematodes, resulting in rapid infestation, yet its overall persistence is lower, indicating that it may be more suitable for controlled environments or specific pest outbreaks.

Overall, the table underscores the importance of selecting the appropriate application method based on persistence and effectiveness, depending on the specific environmental conditions and pest management goals.

## Discussion

This study successfully demonstrates the integration of entomopathogenic nematodes (EPNs) into organic potato farming using an innovative three-stage release technology. The findings reveal significant differences in the efficacy of Steinernema feltiae and Heterorhabditis bacteriophora based on the application methods used—alginate capsules, aqueous suspension, and pest larvae as carriers. Each method offers unique advantages, contributing to sustainable pest management in organic potato cultivation.

Efficacy of the three-stage release technology

The results showed that alginate capsules provided the highest infestation rates and prolonged nematode activity. Specifically, S. feltiae exhibited a 93.6% infestation rate, while H. bacteriophora showed a 91.4% infestation rate under laboratory conditions. This superior performance can be attributed to the waterless environment provided by alginate capsules, which protect nematodes from desiccation and ensure their gradual release into the soil. The low adult emergence rate of 4.6% for S. feltiae further highlights the potential of alginate capsules in reducing pest populations without the need for frequent reapplications.

The aqueous suspension method, although slightly less effective than alginate capsules, still achieved impressive results, with an infestation rate of 92.2% for S. feltiae and 90.1% for H. bacteriophora. This method allowed for a more immediate impact on pest larvae, and its combination of multiple isolates increased virulence. However, the higher adult emergence rate of 5.8% indicates a slightly shorter persistence compared to the capsule method.

The pest larvae application method achieved the highest overall infestation rates, with S. feltiae reaching 95.4% and H. bacteriophora 93.7%. This method allowed for direct interaction between the nematodes and the pests, leading to rapid infestation. However, the higher adult emergence rate (7.4% for S. feltiae and 8.1% for H. bacteriophora) suggests that while this method provides an efficient means of nematode delivery, its persistence may be reduced compared to the other methods.

Epns performance against agrotis segetum pupae

The results demonstrate that all tested EPN species were effective against Agrotis segetum pupae, with S. feltiae showing the highest infestation rate of 81.6% and H. bacteriophora recording 78.4%. These findings align with previous studies highlighting the high efficacy of S. feltiae in managing pupal infestations (Shapiro-Ilan & Lewis, 2014). The higher adult emergence rates in pupae compared to larvae indicate that nematode efficacy decreases slightly as the pest progresses through its life stages. Nevertheless, the use of alginate capsules in managing pupae showed promising results in reducing the overall emergence of adult pests.

Field trials: ecological and economic implications

The field experiments further validated the effectiveness of the three-stage release technology. Alginate capsules proved to be the most environmentally safe method, with infestation rates of 88.2% for larvae and 69.4% for adults. The ability of alginate capsules to maintain nematode viability under dry conditions makes this method particularly well-suited for use in arid and semi-arid regions where organic potato farming is prominent. Additionally, this method’s environmental benefits, including reduced pesticide residues and minimal soil contamination, position it as a key innovation in sustainable agriculture.

The aqueous suspension method, while economically viable and easy to implement, showed slightly lower infestation rates in the field (85.8% for larvae and 64.3% for adults). The rapid dispersal of nematodes in aqueous suspension ensures a faster pest infestation, making this method ideal for immediate pest control interventions. However, the lower persistence compared to alginate capsules suggests that this method may require more frequent applications in field conditions.

The pest larvae application method showed promising results with infestation rates of 83.6% for larvae and 60.8% for adults. While this method offers direct nematode-pest interaction, the rapid decline in nematode performance under field conditions suggests that environmental factors such as soil moisture and temperature may significantly impact its effectiveness.

Ecological and economic efficacy

The three-stage release technology offers a novel and sustainable solution for integrating entomopathogenic nematodes into organic potato farming. The alginate capsules, in particular, provide an innovative approach to maintaining nematode viability and ensuring prolonged pest control. The economic benefits of this technology include reduced pesticide use, lower environmental impact, and improved potato yield due to effective pest management. Furthermore, the ecological benefits of reduced pesticide residues and enhanced soil health underscore the importance of adopting biological control measures in sustainable agriculture.

## Conclusion

The results of this study demonstrated that the three-stage release technology for entomopathogenic nematodes (EPNs) is an innovative, ecologically, and economically effective approach for managing potato pests. The alginate capsules provided prolonged survival and ecological safety of EPNs, ensuring their sustained activity over time. The infestation rates recorded were 93.6% for *Steinernema feltiae* and 91.4% for *Heterorhabditis bacteriophora* under laboratory conditions, indicating the capsules’ ability to create a stable environment, significantly enhancing nematode efficacy.

The aqueous suspension method also proved to be effective, achieving infestation rates of 92.2% for *S. feltiae* and 90.1% for *H. bacteriophora*. The use of multiple isolates combined in the suspension contributed to increased infestation rates. While this method resulted in a slightly higher rate of adult emergence compared to alginate capsules, it was both practically feasible and cost-effective.

The pest larvae-based application exhibited the highest infestation rates, with *S. feltiae* reaching 95.4% and *H. bacteriophora* achieving 93.7%. This method ensured direct contact between nematodes and pests, thereby enhancing overall efficacy. This technology, particularly through the use of alginate capsules and aqueous suspension, ensures reduced pesticide residues and improved environmental safety while also conserving water resources.

The statistical analysis, conducted through ANOVA and LSD tests (α = 0.05), confirmed the scientific validity of the results, with significant impacts observed in pest infestation rates (p<0.05). These findings affirm that EPNs can be an effective biocontrol strategy for managing potato pests in organic agriculture.

The comparison with previous studies showed similarities in EPNs’ persistence and efficacy; however, the three-stage technology introduced in this study offers distinct innovations. For example, the prolonged survival rates observed with alginate capsules and the increased virulence achieved through the aqueous suspension align with earlier studies by Shapiro-Ilan & Lewis (2014). This integrated release strategy thus represents an effective and promising approach for sustainable potato farming.

Practically, the three-stage release technology can be recommended for use in organic potato cultivation to control pests. Alginates are ideal for dry conditions, while the aqueous suspension and pest larvae-based applications are suitable for broader field use due to their cost-effectiveness and efficiency.

## Author Contributions

Azimjon Anorbayev conceptualized the study and conducted laboratory experiments. Nafosat Kurbonova developed the methodology, organized the field trials, and analyzed the data. Nigora Tillyakhodjaeva edited the manuscript, summarized the findings, and contributed to writing the scientific article. All authors reviewed and contributed to the editing process.

This article is based on collaborative research on implementing an innovative three-stage EPN release technology in organic potato farming, with each author actively contributing to the design, execution, analysis, and writing of the study.

## Supporting information

Laboratory protocols

