## Supplementary material for "Integrating entomopathogenic nematodes into sustainable organic potato farming through a three-stage release technology": Laboratory protocols

***Research Laboratory Protocols***

**1. Alginate Capsule Preparation Protocol**

**Objective:**

To develop waterless alginate capsules for effective and ecological distribution of entomopathogenic nematodes (EPN).

**Materials:**

- 2% sodium alginate solution
- Calcium chloride solution (0.2 M)
- Entomopathogenic nematodes containing 500 infective juveniles (IJs)
- Sterile distilled water
- Petri dishes (9 cm in diameter)
- Whatman No.1 filter paper
- Micropipette
- Parafilm

**Steps:**

1. **Preparation of alginate solution**:
   - Prepare a 2% sodium alginate solution. Carefully add 500 infective juveniles (IJs) to the solution and mix thoroughly.
2. **Capsulation process**:
   - Using a micropipette, drop the sodium alginate solution into the 0.2 M calcium chloride solution, forming capsules.
   - Allow the capsules to remain in the calcium chloride solution for 5-10 minutes to solidify.
3. **Capsule preparation**:
   - Remove the capsules from the calcium chloride solution, rinse them with sterile distilled water, and dry them.
   - Place the capsules into Petri dishes lined with Whatman No.1 filter paper, adding 1.5 ml of sterile distilled water to maintain moisture.
   - Seal the dishes with Parafilm and incubate at 18 ± 1°C for 48 hours.
4. **Capsule evaluation**:
   - After incubation, check the capsules to ensure they retain EPN viability and infectivity for further experiments.

**Outcome:**

Alginate capsules enable continuous EPN distribution under water-deficient conditions, ensuring prolonged nematode activity.

**2. Aqueous Suspension Preparation Protocol**

**Objective:**

To achieve broader efficacy by combining multiple EPN isolates in an aqueous suspension.

**Materials:**

- 0.1 ml aqueous suspension containing approximately 100 infective juveniles (IJs)
- Petri dishes (9 cm)
- Whatman No.1 filter paper
- Sterile distilled water
- Micropipette
- Parafilm

**Steps:**

1. **Preparation of the aqueous suspension**:
   - Mix 100 infective juveniles (IJs) with sterile distilled water to create a 0.1 ml suspension.
   - Add 0.1 ml of the suspension onto the filter paper in Petri dishes.
2. **Incubation process**:
   - Place five *Agrotis segetum* larvae in each Petri dish and incubate at 18 ± 1°C for 4 days.
   - Seal the dishes with Parafilm to maintain humidity throughout the incubation period.
3. **Maintaining moisture in the suspension**:
   - Regularly monitor the moisture level of the filter paper throughout the experiment, adding sterile water as needed to maintain optimal conditions for EPN activity.

**Outcome:**

The combined aqueous suspension of multiple EPN isolates ensures faster pest infection and enhances overall efficacy.

**3. EPN Distribution via Larvae Protocol**

**Objective:**

To distribute EPNs directly to pests using *Agrotis segetum* larvae as carriers.

**Materials:**

- *Agrotis segetum* larvae (5-7 days old)
- Entomopathogenic nematodes containing 500 infective juveniles (IJs)
- Petri dishes
- Micropipette
- Distilled water

**Steps:**

1. **Preparation of larvae**:
   - Place 5-7 day-old *Agrotis segetum* larvae into Petri dishes, as they exhibit the highest susceptibility.
   - Apply 500 infective juveniles (IJs) to each larva and allow them to interact for 24 hours.
2. **Distribution through larvae**:
   - After 24 hours of interaction, release the infected larvae into the field or lab environment. This approach directly transfers nematodes to the target pests.
3. **Monitoring larval infection**:
   - Continuously observe the infection rate of the larvae and measure the speed and extent of pest infection.

**Outcome:**

Larval-based distribution provides an effective means for directly delivering EPNs to pests, resulting in high infection rates.
